# Peripheral Na_V_1.7 modulation reveals divergent peripheral and central adaptations in persistent trigeminal pain

**DOI:** 10.64898/2026.09.16.752059

**Authors:** Heather N. Allen, Naomi K. Grabus, Rajesh Khanna

**Author notes:** Corresponding author: Rajesh Khanna, PhD, Mailing address: 1149 Newell Dr, L4-183, Gainesville, FL, 32610, USA. authors contributed equally to this work (Heather N. Allen, Naomi K. Grabus).

## Abstract

Trigeminal pain produces persistent facial hypersensitivity that is difficult to treat. Ion channels contribute to the pathology of trigeminal pain, but the consequences of their modulation remain poorly understood. We previously demonstrated that voltage-gated sodium channel Na_v_1.7 is a therapeutic target that can be indirectly modulated via interaction with collapsin mediated response protein 2 (CRMP2) to reduce membrane expression of Na_v_1.7 and trigeminal ganglion neuronal excitability. We used the Foramen Rotundum Inflammatory Constriction of the Trigeminal InfraOrbital Nerve (FRICT-ION) model of trigeminal neuropathic pain to characterize behavioral and supraspinal effects of indirectly modulating the peripheral CRMP2-Na_v_1.7 interaction via our novel Compound 194 (C194). FRICT-ION produced mechanical and cold allodynia, which were reduced by C194, but had no effect on non-evoked affective behaviors. Despite behavioral improvement, C194 did not normalize heightened glutamatergic activity in the parabrachial nucleus (PBN), revealing a dissociation between behavioral analgesia and activity of a central pain circuit. These findings extend mechanistic studies of C194 in the trigeminal system and demonstrate efficacy of CRMP2-Na_v_1.7 disruption in modulating facial pain-like behavior. These results support the CRMP2-Na_v_1.7 axis as a promising therapeutic target while highlighting the need to define how peripheral and central adaptations interact to shape persistent facial pain.

## Introduction

Facial neuropathic pain encompasses a heterogeneous group of disorders affecting the trigeminal somatosensory system, including conditions like posttraumatic trigeminal neuropathic pain (PTNP), trigeminal neuralgia (TN), and other persistent orofacial pain conditions(1). These conditions can produce severe, often incapacitating pain that interferes with routine activities like eating, speaking, and touching the face(2). TN represents one of the most severe and clinically well-characterized forms of trigeminal pain and is classically defined by recurrent, unilateral, electric shock-like pain provoked by innocuous stimuli(3). PTNP often develops following injury to a trigeminal nerve, leading to persistent pain(4). The pathophysiology of trigeminal pain is complex and incompletely understood, leading diagnosis difficulty, incomplete treatment, and patient frustration and suffering(3,5). Although primary TN is often associated with neurovascular compression and focal demyelination of the trigeminal root entry zone, traumatic injury, inflammation, and peripheral nerve dysfunction contribute to other forms of trigeminal neuropathic pain(3,5). Further, the burden of trigeminal pain extends beyond sensory symptoms; patients with trigeminal pain exhibit increased rates of anxiety and depression compared to the general population as well as other chronic pain populations(6–8). Although trigeminal pain conditions exhibit differences in clinical presentation and etiology, they share pathological features of trigeminal nerve dysfunction and aberrant neuron excitability,(3,9,10) suggesting that structural nerve pathology alone does not fully explain the development or persistence of trigeminal pain. These diverse presentations support the role for maladaptive changes in the molecular mechanisms governing neuronal excitability in trigeminal pain. There is a significant need to better understand the mechanisms underlying trigeminal pain to develop improved therapeutics.

Voltage-gated sodium (Na_v_) channels are central regulators of neuronal excitability and are essential for pain signaling(11–13). Dysregulation of Na_v_ channels can promote aberrant spontaneous firing, lower neuronal firing threshold, and facilitate the transmission of nociceptive signals(14). Na_v_ channels remain a top target for pharmacological treatment of trigeminal pain(5,15): Carbamazepine (CBZ) and oxcarbamazepine, the first line treatment for patients with TN, broadly target sodium channels, but their nonselective activity is associated with adverse effects that significantly affect patient adherence to treatment(3,5,16). Therapeutics directed at more selective targets may provide effective analgesia while reducing off-target effects.

Multiple Na_v_ channel subtypes, including Na_v_1.3, Na_v_1.7, and Na_v_1.8, are expressed in trigeminal ganglion (TG) neurons and have been implicated in trigeminal pain (17,18). Emerging evidence indicates that voltage-gated sodium channel Na_v_1.7, encoded by *Scn9a,* represents a particularly compelling therapeutic target due to its role in peripheral neuron excitability and function in human pain(12,13). Data from human trigeminal disease further supports a role for Na_v_1.7 in pathological facial pain: various mutations in *Scn9a* have been identified across studies in patients with trigeminal pain(18,19) and changes in Na_v_1.7 function have been identified in persistent pain following trigeminal injury(20). However, despite accumulating evidence for Na_v_1.7 in pain conditions, therapeutic translation has proven difficult(21). Vixotrigine, a selective Na_v_1.7 blocker used in a phase 2a clinical trial in patients with TN, reduced the number and severity of paroxysms and daily pain scores compared to placebo controls despite the endpoint not reaching statistical significance (22,23). While not reaching statistical significance for the primary endpoint, the trial provided proof of concept for Na_v_1.7-directed therapy in TN and support continued investigation of strategies to modulate Na_v_1.7 in trigeminal pain.

Preclinical studies provide additional evidence that Na_v_1.7 contributes to trigeminal pain and reducing its function can alleviate pain-related behaviors. Rats with temporomandibular joint inflammation exhibit greater Na_v_1.7 expression and sodium currents in TG neurons(24,25), suggesting that inflammatory signaling can enhance Na_v_1.7-dependent excitability in the trigeminal system. Selective inhibition of Na_v_1.7 attenuates TG activity after pulpitis-induced inflammatory orofacial pain(26), supporting a causal role of Na_v_1.7 in facial pain. In models of trigeminal nerve injury, modulation of signaling pathways that regulate Na_v_1.7 has also been associated with reductions in mechanical allodynia. Botulinum toxin type A decreases infraorbital nerve-injury-induced pain-like behaviors via reduction of calcitonin gene-related peptide (CGRP)-dependent signaling, which was found to induce extracellular regulated kinase (ERK)-dependent reduction of Na_v_1.7 membrane localization and sodium current density(27). Similarly, pituitary adenylyl cyclase activating peptide (PACAP)-mediated attenuation of trigeminal neuropathic pain has also been linked to ERK-dependent modulation of Na_v_1.7(28). Therefore, regulation of Na_v_1.7 function via upstream signaling pathways may represent a common mechanism through which diverse interventions alleviate trigeminal pain.

Therapeutics aimed at indirect targeting of Na_v_1.7 via regulatory proteins that control expression, trafficking, or membrane activity provide an opportunity to selectively decrease pathological Na_v_1.7 activity while preserving physiological sodium channel function(29–31). One such approach involves collapsin response mediator protein 2 (CRMP2), a cytosolic protein that regulates the trafficking and functional expression of Na_v_1.7 at the plasma membrane(32). CRMP2 interacts with Na_v_1.7 and promotes membrane localization of the channel: this interaction is regulated by SUMOylation of CRMP2, which contributes to maintenance of functional Na_v_1.7 expression at the membrane(32). DeSUMOylation of CRMP2 promotes Nav1.7 internalization, reducing trafficking and sodium currents(32). Compound 194 (C194) was developed to disrupt the CRMP2-Na_v_1.7 regulatory axis and has been extensively characterized as an indirect modulator of Na_v_1.7. C194 reduces Na_v_1.7-dependent sodium currents, decreases sensory neuron excitability, and alleviates pain-related behaviors across multiple preclinical models of neuropathic and inflammatory pain(32–37). We previously demonstrated that the mechanisms established for C194 in the dorsal root ganglia (DRG) extend to the trigeminal system as well, reducing Na_v_1.7 expression at the membrane and decreasing excitability of TG neurons while also attenuating pain-like behaviors associated with chronic constriction injury of the infraorbital nerve (CCI-ION)(35).

Here, we use C194 in the FRICT-ION (Foramen Rotundum Inflammatory Constriction of the Trigeminal InfraOrbital Nerve) model to further explore the role of Na_v_1.7 modulation in trigeminal pain. FRICT-ION is a minimally invasive mouse model used to investigate chronic trigeminal neuropathic pain that produces persistent facial mechanical allodynia(38,39). Although FRICT-ION does not reproduce the full clinical phenotype or etiology of human TN, it provides a complementary model of persistent trigeminal pain to investigate the consequences of chronic facial neuropathic pain over extended periods. Building on our previous work establishing the efficacy and mechanism of C194 on peripheral pain models, we integrate *in situ* hybridization, *in vivo* calcium imaging, and behavior to define how indirect Na_v_1.7 modulation can be used to relieve trigeminal neuropathic pain.

## Results

Wildtype male and female mice underwent FRICT-ION surgery to induce trigeminal facial pain or a sham surgery before being tested for pain like behaviors (**Figure 1A**). Compared to sham animals, FRICT-ION mice developed both robust mechanical allodynia and cold hypersensitivity starting seven days post-surgery, confirming successful induction of a trigeminal neuropathic pain phenotype (**Figure 1B-C).** To determine whether FRICT-ION reflects a clinically relevant model, we evaluated the effects of carbamazepine (CBZ), a first-line therapy for TN patients. Following the development of pain-like behaviors, animals treated with CBZ (60 mg/kg, p.o.) displayed significantly reduced mechanical and cold hypersensitivity (**Figure 1D-E**), supporting the translational relevance of the FRICT-ION model and its responsiveness to standard-of-care therapy. Treatment with vehicle had no effect on FRICT-ION-induced hypersensitivity (**Figure 1D-E**).

**Figure 1.**
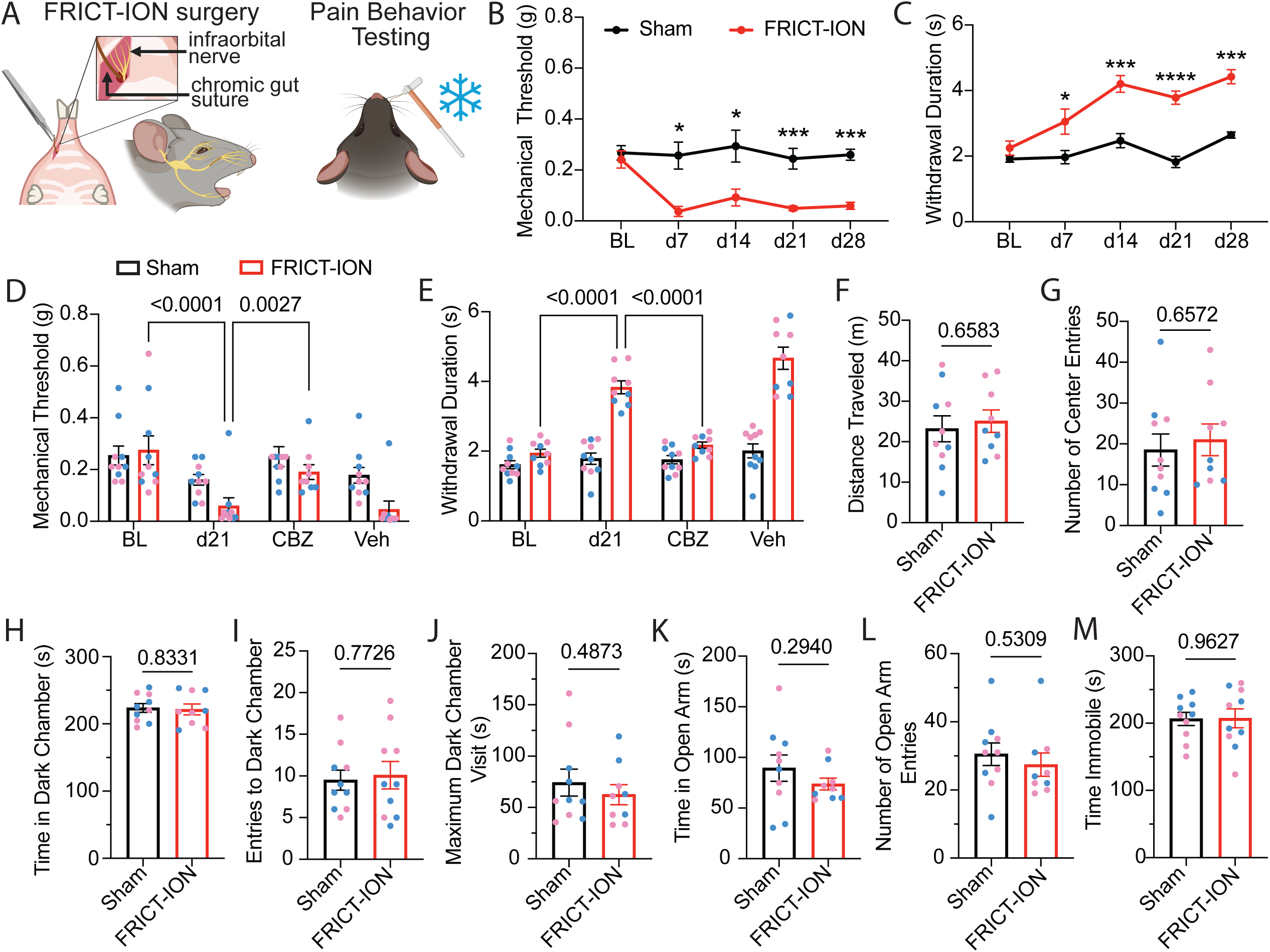
FRICT-ION induces persistent, evoked pain-like behaviors without affective comorbidities. (A) Schematic of FRICT-ION surgery and subsequent behavioral testing. (B) Mechanical withdrawal thresholds measured at baseline and following 28 days post FRICT-ION or sham surgery (Two-way ANOVA, effect of time/drug, p=0.0173, effect of surgery, p<0.0001, effect of interaction, p=0.0436, Holm-Šidák multiple comparisons, n=9-10). (C) Cold withdrawal duration measured at baseline and following 28 days post FRICT-ION or sham surgery (Two-way ANOVA, effect of time/drug, p=<0.0001, effect of surgery, p=<0.0001, effect of interaction, p=0.0019, Holm-Šidák multiple comparisons, n=9-10). (D) Mechanical withdrawal thresholds at baseline, post FRICT-ION or sham surgery, and following carbamazepine (CBZ) or vehicle treatment (Two-way ANOVA, effect of time/drug, p<0.0001, effect of surgery, p=0.0227, effect of interaction, p=0.0492, Tukey’s multiple comparisons, n=10) (E) Cold withdrawal duration measured at baseline, post FRICT-ION or sham surgery, and following treatment with CBZ or vehicle (Two-way ANOVA, effect of time/drug, p<0.0001, effect of surgery, p<0.0001, effect of interaction, p<0.0001, Tukey’s multiple comparisons, n=10). (F) Total distance traveled in the open-field test in sham and FRICT-ION mice. (G) Number of entries in the center of the open-field test apparatus in sham and FRICT-ION mice. (H) Total time spent in the dark chamber of the light-dark box test apparatus in sham and FRICT-ION mice. (I) Number of entries into the dark chamber of the light-dark box test apparatus in sham and FRICT-ION mice. (J) Maximum amount of time spent in the dark chamber of the light-dark box test apparatus in sham and FRICT-ION mice. (K) Total time spent in the open arm of the elevated-plus maze test apparatus in sham and FRICT-ION mice. (L) Number of entries into the open arm of the elevated-plus maze test apparatus in sham and FRICT-ION mice. (M) Total time spent immobile in the forced-swim test in sham and FRICT-ION mice, (F-M unpaired Welch’s t test, n=9-10). Pink data points represent female subjects; blue data points represent male subjects. *p<0.05, **p<0.01, ***p<0.001, ****p<0.0001.

To assess if the FRICT-ION model induces affective comorbidities associated with persistent trigeminal pain, mice underwent behavioral testing 10 weeks following surgery. Importantly, locomotor activity remained unchanged between sham and FRICT-ION mice, indicating that the observed behavioral responses were not due to changes in mobility or motor impairment **(Figure 1F)**. No significant differences were observed between sham and FRICT-ION mice in assays for anxiety- or depression-like behaviors **(Figure 1G-M)**. Despite the presence of robust evoked pain-like behaviors, the FRICT-ION model does not produce affective pain-like behaviors in our hands at ten weeks post-surgery.

Dysregulation of Na_v_1.7 channels is associated with many chronic pain conditions, including trigeminal neuropathic pain. To test whether Na_v_1.7 channels contribute to facial pain in the FRICT-ION model, we administered compound 194 (C194), a small molecule that targets CRMP2-mediated regulation of Na_v_1.7(32), to sham and FRICT-ION mice and assessed facial mechanical and cold allodynia (**Figure 2A**). Intraperitoneal administration of C194 (10 mg/kg) significantly increased the mechanical withdrawal threshold of mice with FRICT-ION but had no effect in sham animals (**Figure 2B**). Similarly, C194 alleviated the cold hypersensitivity observed in FRICT-ION mice 21 days post-surgery (**Figure 2C**). We also tested the efficacy of C194 when administered intranasally to assess translational modes of treatment (**Figure 2D**). Intranasal C194 (1 mg/mouse) effectively reversed FRICT-ION-induced mechanical and cold allodynia (**Figure 2E-F**), demonstrating therapeutic potential.

**Figure 2.**
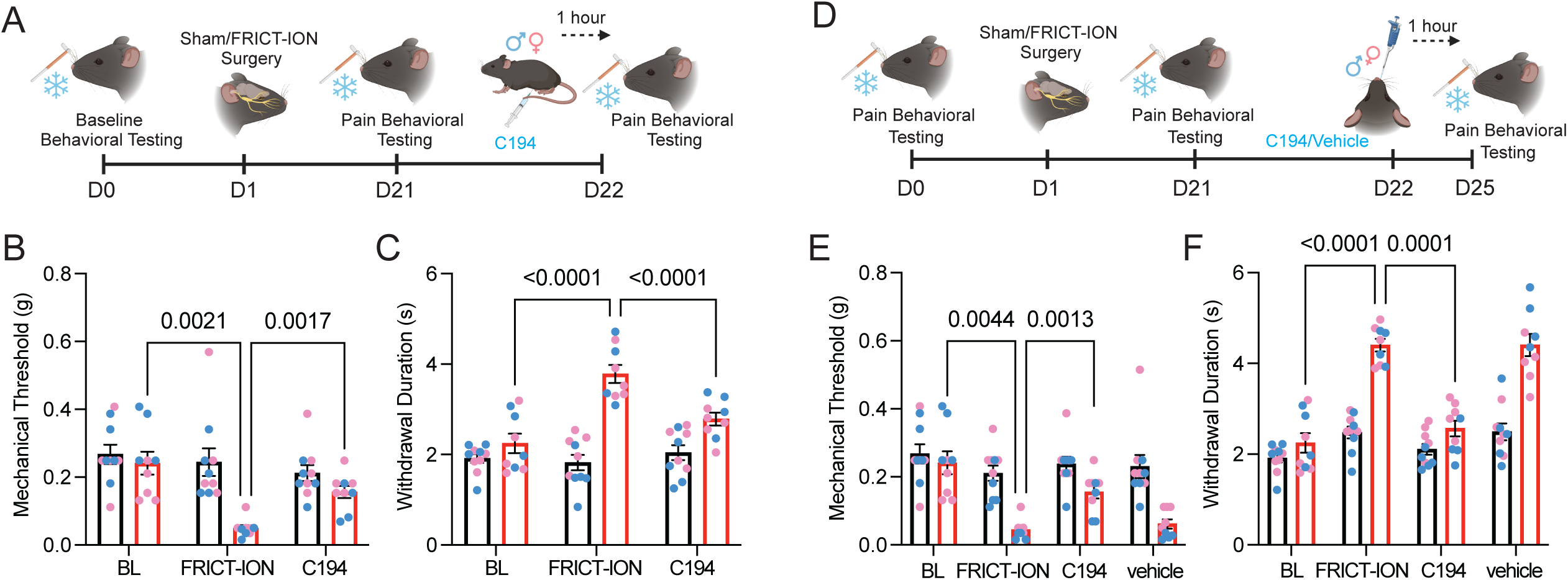
Compound 194 attenuates mechanical and cold hypersensitivity in FRICT-ION mice. (A) Schematic of the experimental timeline of systemic C194 in Sham/FRICT-ION animals (B) Mechanical withdrawal thresholds measured at baseline, post FRICT-ION or sham surgery, and following intraperitoneal treatment with C194 (Two-way ANOVA, effect of time/drug, p=0.0115, effect of surgery, p=0.0002, effect of interaction, p=0.0348, Tukey’s multiple comparisons, n=9-10). (C) Cold withdrawal duration measured at baseline, post FRICT-ION or sham surgery, and following intraperitoneal treatment with C194 (Two-way ANOVA, effect of time/drug, p<0.0001, effect of surgery, p<0.0001, effect of interaction, p<0.0001, Tukey’s multiple comparisons, n=9-10). (D) Schematic of the experimental timeline of intranasal C194 in Sham/FRICT-ION animals. (E) Mechanical withdrawal thresholds measured at baseline, post FRICT-ION or sham surgery, and following intranasal treatment with C194 (Two-way ANOVA, effect of time/drug, p=0.0005, effect of surgery, p<0.0001, effect of interaction, p=0.0342, Tukey’s multiple comparisons, n=9-10). (F) Cold withdrawal duration measured at baseline, post FRICT-ION or sham surgery, and following intranasal treatment with C194 (Two-way ANOVA, effect of time/drug, p<0.0001, effect of surgery, p<0.0001, effect of interaction, p<0.0001, Tukey’s multiple comparisons, n=9-10). Pink data points represent female subjects; blue data points represent male subjects.

Given that C194 reduces pain-like behaviors through modulation of CRMP2-Nav1.7 interactions, we hypothesized that genetic disruption of this regulatory pathway would similarly alter the progression of TN-related pain-like behaviors. Mice with a genetic deletion of the CRMP2 regulatory sequence (ΔCRS) (40) underwent FRICT-ION or sham surgery and repeated mechanical and cold sensitivity testing to determine if CRMP2 regulation of Na_v_1.7 is essential to the development of trigeminal neuropathic pain (**Figure 3A**). Deletion of the CRS sequence did not affect expression of *Scn9a* or *Dpysl2* (encoding CRMP2) in the TG (**Figure 3B-C**). Wildtype mice that received FRICT-ION surgery developed persistent facial mechanical and cold hypersensitivity compared to wildtype mice that received sham surgery starting on day 14 post-surgery that lasted for at least 84 days (**Figure 3D-E**). However, ΔCRS mice that underwent FRICT-ION surgery did not develop the same mechanical and cold hypersensitivity as wildtype FRICT-ION mice. Sensitivity of ΔCRS FRICT-ION mice was not significantly different from wildtype sham or ΔCRS sham, suggesting that the CRMP2-Na_v_1.7 interaction is necessary for the development of trigeminal neuropathic pain.

**Figure 3.**
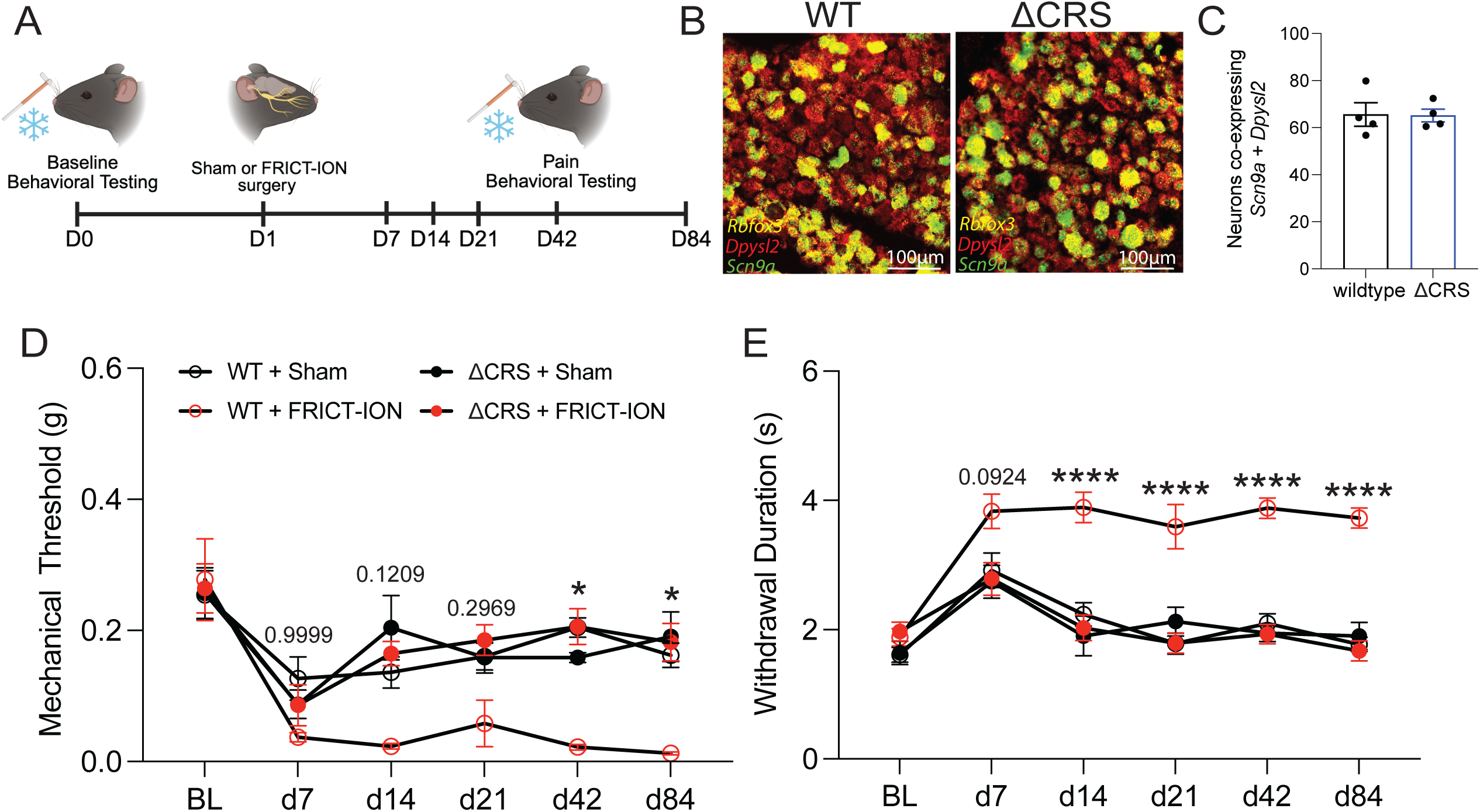
CRMP2 regulatory sequence mutation reduces mechanical and cold hypersensitivity. (A) Schematic of the experimental timeline. (B) Representative *in situ* hybridization images demonstrating neuronal expression of *Dpysl2* and *Scn9a* in the trigeminal ganglion of wildtype and ΔCRS mice. (C) Quantification of neurons co-expressing *Dpysl2* and *Scn9a* in the trigeminal ganglion of wildtype and ΔCRS mice (Unpaired t-test, 0.9377, n=4 animals, each data point represents the average of 3-4 TG sections). (D) Mechanical withdrawal thresholds measured at baseline and then 7, 14, 21, 42, 84 days following FRICT-ION or sham surgery in wild type and CRS mutant mice (Three-way ANOVA, effect of time x genotype x surgery, p=0.0249, Tukey’s multiple comparisons, WT FRICT-ION vs. CRS FRICT-ION, n=9-11). (E) Cold withdrawal duration measured at baseline and then 7, 14, 21, 42, 84 days following FRICT-ION or sham surgery in wild type and CRS mutant mice (Three-way ANOVA, effect of time x genotype x surgery, p=0.0002, Tukey’s multiple comparisons, WT FRICT-ION vs. CRS FRICT-ION, n=9-11). *p<0.05, **p<0.01, ***p<0.001, ****p<0.0001.

While nociceptive signals are generated in the periphery, the central nervous system is imperative for creating the perception of pain and is highly involved in the modulation of persistent pain(41,42). The parabrachial nucleus (PBN) is an essential relay for the transmission of nociceptive signals from the trigeminal ganglia to the brain, receiving both direct TG input as well as input via spinal trigeminal nucleus(43–45). Although this circuitry has largely been defined in rodents, ultra-high-field diffusion MRI has recently delineated a direct trigeminal–lateral parabrachial–central amygdala tract in humans, providing in vivo confirmation that trigeminal nociceptive information reaches the PBN in the human brain(46). Additionally, changes in the PBN have been observed in humans during pain(47–50), underscoring its importance as a key relay structure for supraspinal pain signals. To assess activity of the PBN in trigeminal pain, we labeled immediate early gene *Fos* in the PBN of sham and FRICT-ION mice (**Figure 4A**). FRICT-ION mice displayed higher *Fos* expression in the PBN compared to sham controls (**Figure 4B**), demonstrating increased activation of PBN neurons in the context of neuropathic facial pain. Interestingly, there was no significant difference between the number of *Fos* positive cells in the left and right PBN, despite the injury being on the right side of the face (**Figure 4C)**, suggesting equal activation of the bilateral PBN in the context of trigeminal neuropathic facial pain.

**Figure 4.**
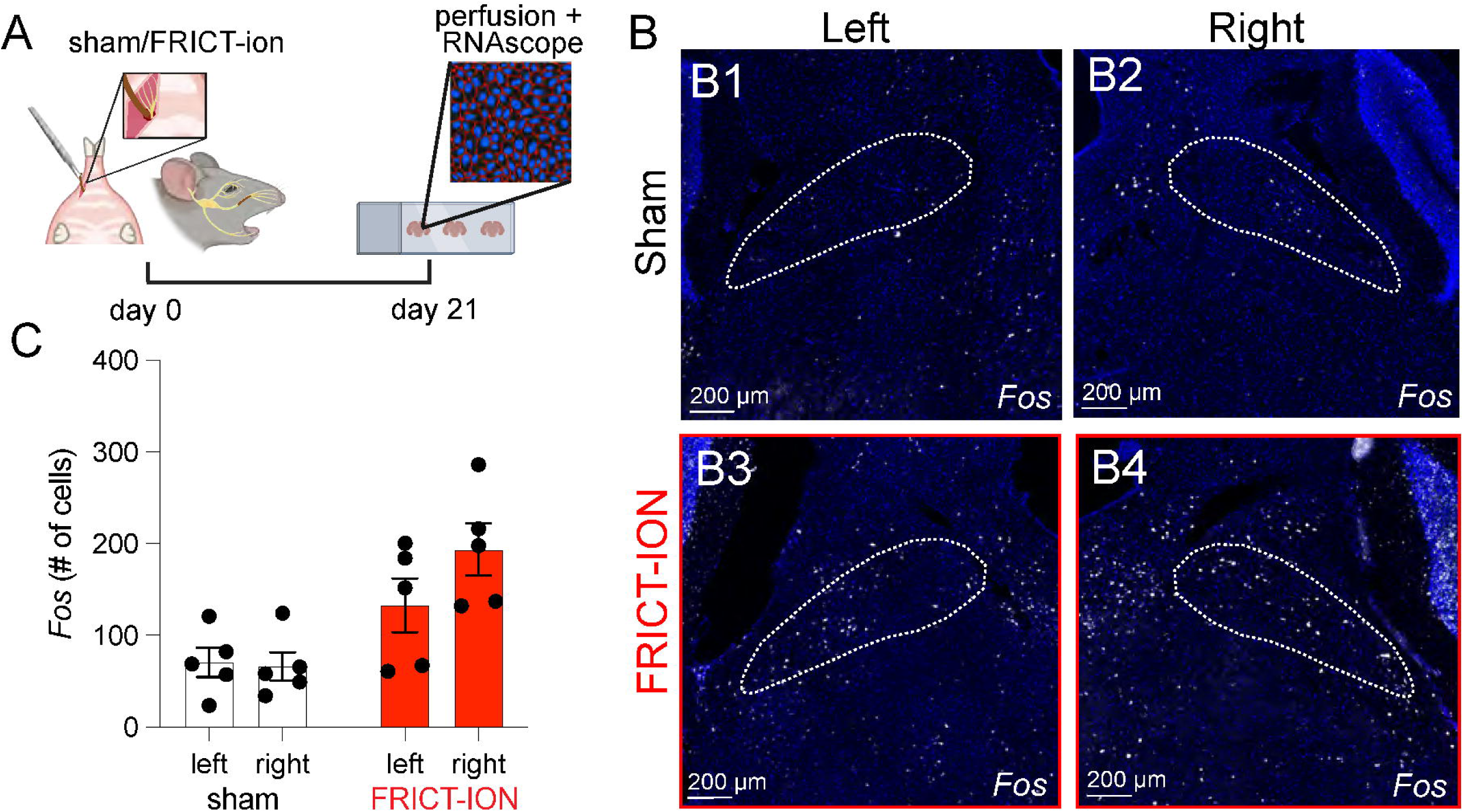
FRICT-ION increases neuronal activation in the parabrachial nucleus. (**A**) Experimental schematic for *Fos* in the PBN. (**B**) Representative images for *Fos* in the left (**B1**) and right (**B2**) PBN of sham animals compared to left (**B3**) and right (**B4**) PBN of FRICT-ION animals. (**C**) Quantification of *Fos* positive cells in the PBN of sham and FRICT-ION animals (Two-way RM ANOVA, effect of injury p=0.0008, effect of hemisphere p=0.2387, effect of interaction p=0.1791, n=5, each data point represents the average of 2-4 sections from one animal).

To assess activation changes in the PBN *in vivo*, calcium imaging in glutamatergic PBN neurons was recorded using fiber photometry before and after FRICT-ION surgery (**Figure 5A-B**). PBN neuronal activity trended toward increasing in response to light (**Figure 5C-D**) and heavy (**Figure 5I-J**) mechanical stimulation of the face and significantly increased in response to cold stimulation (**Figure 5O-P**) after FRICT-ION, suggesting enhanced supraspinal nociceptive signaling following trigeminal nerve injury. Pharmacological administration of analgesics has been shown to diminish stimulus-evoked PBN responses in the context of body pain-induced hypersensitivity in rodents(33,34,51–53). Hence, we administered C194 to test if the analgesic effects observed in behavior would be represented in the nociceptive signaling pathway in the brain. Interestingly, neither C194 (10 mg/kg) nor carbamazepine (60 mg/kg) had any effect on responses of glutamatergic PBN neurons to mechanical and cold stimulation (**Figure 5E-T**). Although FRICT-ION increased responses of PBN neurons to evoked stimuli (**Figure 5U-W**), pharmacological therapeutics did not alter PBN activity further despite reducing behavioral readouts (**Figures 1-2**), indicating that the analgesic effects of CBZ and C194 do not require normalization of parabrachial signaling to be effective.

**Figure 5.**
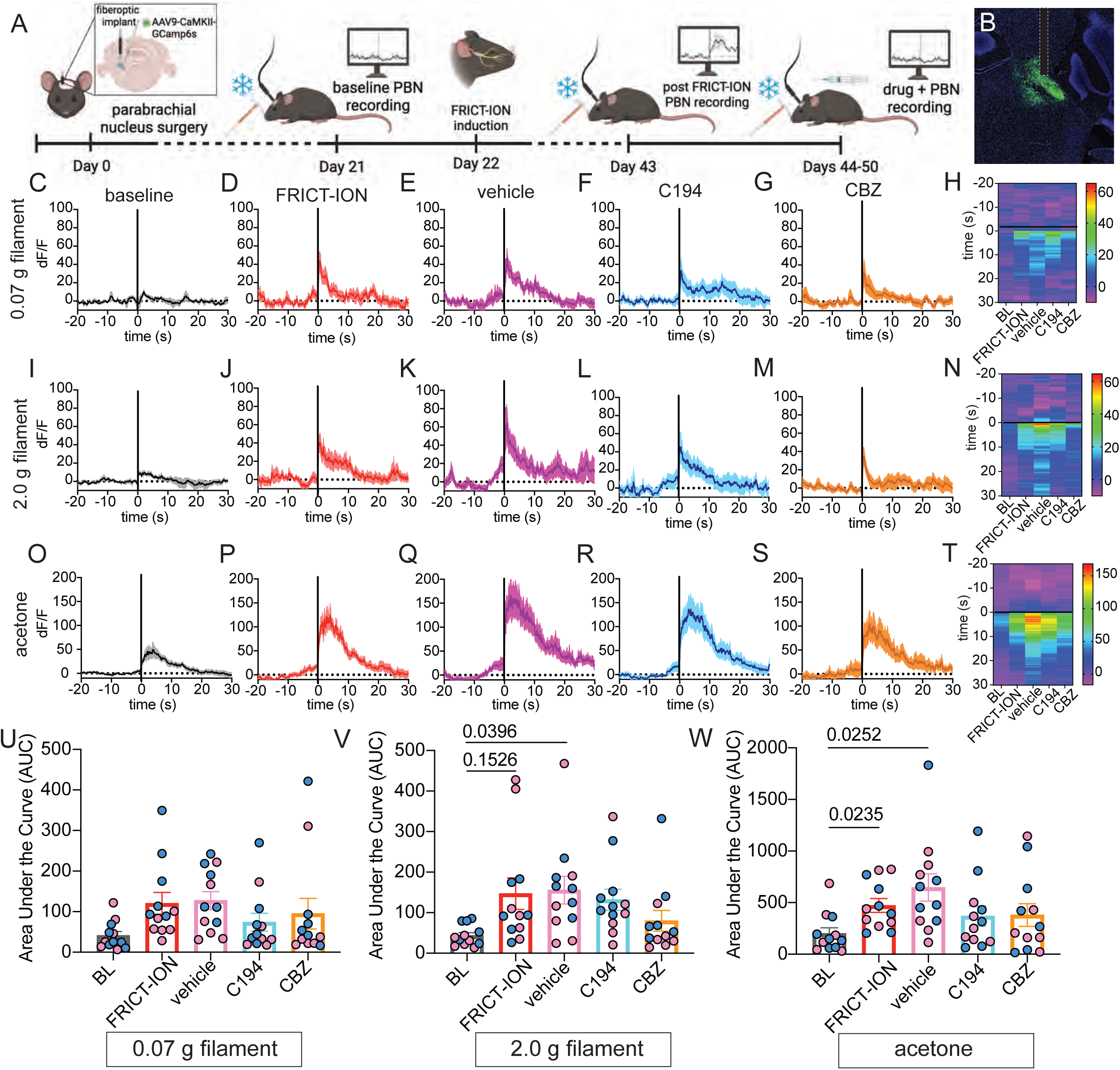
FRICT-ION increases stimulus evoked PBN neuronal activity. (A) Schematic of the experimental timeline for *in vivo* fiber photometry at baseline, following FRICT-ION surgery, and treatment with C194, CBZ, and vehicle. (B) Representative viral expression of GCamp6 in glutamatergic neurons in the PBN. (C) Averaged calcium transients in response to 0.07 g von Frey filament at baseline, (D) after FRICT-ION surgery, (E) vehicle, (F) C194, (G) and CBZ. (H) Heat map depicting the mean responses across animals at baseline, following FRICT-ION surgery, and treatment with vehicle, C194, and CBZ. (I) Averaged calcium transients in response to 2.0 g von Frey filament at baseline, (J) after FRICT-ION surgery, (K) vehicle, (L) C194, (M) and CBZ. (N) Heat map depicting the mean responses across animals at baseline, following FRICT-ION surgery, and treatment with vehicle, C194, and CBZ. (O) Averaged calcium transients in response to 10 μL droplet of acetone at baseline, (P) after FRICT-ION surgery, (Q) vehicle, (R) C194, (S) and CBZ. (T) Heat map depicting the mean responses across animals at baseline, following FRICT-ION surgery, and treatment with vehicle, C194, and CBZ. (U) Area under the curve (AUC) analysis of responses recorded during the five-second period following 0.07 g mechanical stimulation (One-way ANOVA, p=0.0553, n=12). (V) Area under the curve (AUC) analysis of responses recorded during the five-second period following 2.0 g mechanical stimulation (One-way ANOVA, p=0.0348, Tukey’s multiple comparisons, n=12). (W) Area under the curve (AUC) analysis of responses recorded during the five-second period following acetone cold stimulation (One-way ANOVA, p=0.0035, Tukey’s multiple comparisons, n=12). The timing of stimulus application is denoted by a solid black line. Solid traces indicate mean responses, with shaded regions representing SEM. Each point in the AUC plots corresponds to an individual animal. Pink data points represent female subjects; blue data points represent male subjects.

## Discussion

The present study provides further evidence that indirect modulation of Na_v_1.7 through the CRMP2 pathway alleviates trigeminal neuropathic pain in mice. FRICT-ION produced robust and persistent facial mechanical and cold allodynia that were responsive to carbamazepine, supporting the sensitivity of this model to clinically established treatment. C194 similarly reduced established mechanical and cold hypersensitivity following FRICT-ION, with efficacy observed following both intraperitoneal and intranasal administration. Together with our previous work which establishes the molecular mechanisms underlying C194 activity in the trigeminal system(35), these findings support the Na_v_1.7-CRMP2 regulatory axis as a therapeutically relevant mechanism across multiple dimensions of trigeminal neuropathic pain.

Data from mice lacking the CRMP2 regulatory sequence in Na_v_1.7 further support the importance of this regulatory mechanism. The fact that ΔCRS mice did not develop the same mechanical or cold allodynia following FRICT-ION surgery as wildtype mice complements the pharmacological effects of C194 and provides independent evidence that the CRMP2-Na_v_1.7 interaction contributes to the development of trigeminal neuropathic pain. Notably, the deletion of the CRS prevents pain from developing after FRICT-ION surgery, whereas C194 administered after the onset of neuropathic pain reverses established facial allodynia. Together, these data suggest the CRMP2-Na_v_1.7 interaction is essential to both the development and maintenance of trigeminal hypersensitivity, suggesting a therapeutically accessible mechanism after pain has become established.

The efficacy of C194 and influence of Na_v_1.7 on trigeminal pain observed here is particularly interesting when comparing to patient data: Na_v_1.7 expression is reported to be reduced in TN patients(17). Total Na_v_1.7 expression, however, is not equal to the functionally available Na_v_1.7 at the membrane. C194 acts by disrupting the CRMP2-dependent trafficking of Na_v_1.7, reducing the availability of functional channels at the membrane without affecting expression of *Scn9a*(54). It is possible that pathological pain depends on the activity or localization of a relatively small population of functional Na_v_1.7, rather than channel abundance. A decrease in Na_v_1.7 expression could also be the result of compensatory remodeling after pain that varies across disease stages, etiology, or neuronal population. Functional regulation of Na_v_1.7 may be more relevant than expression alone and therefore a more relevant therapeutic target.

C194 had a slightly greater effect on cold hypersensitivity then on mechanical sensitivity, suggesting that these behaviors may have partially distinct mechanisms. Mechanical allodynia and cold allodynia can arise from different populations of primary sensory neurons(55,56), and their dependence on sodium channels varies with the nature of the nerve injury(57). Mouse models of neuropathic pain also reveal recruitment of large diameter, normally silent sensory neurons that become responsive to cold(58). Cold sensitivity is likely a combination of altered peripheral excitability and recruitment of sensory populations distinct from those driving mechanical sensitivity. C194 completely reversed cold allodynia of FRICT-ION mice, while mechanical allodynia was alleviated but did not return to baseline levels, possibly representing preferential modulation of a Na_v_1.7 population contributing to pathological cold signaling. Consistent with this hypothesis, human neuroimaging studies reveal differential activation of brainstem and supraspinal areas in response to mechanical and cold stimuli in patients with trigeminal neuropathic pain(59), suggesting stimulus-specific signaling pathways in the central nervous system. Further definition of the sensory neurons affected by C194 will be important for further disentangling specific pain modalities.

Calcium imaging of glutamatergic PBN neurons reveals dissociation between behavioral analgesia and activity of the central nervous system. FRICT-ION significantly increased cold-evoked PBN activity and trended toward an increase in mechanical-evoked PBN activity, yet neither C194 nor CBZ normalized these responses, despite alleviating behavioral hypersensitivity to the same stimuli. Rodriguez et al. identify a direct projection from the TG to PBN and demonstrate that this pathway is preferentially activated by facial pain(43). Persistent PBN activity in the presence of C194 and CBZ may indicate continued recruitment of a circuit that is important for facial pain but not required for behavioral improvement: C194 may be alleviating pain-like behaviors through signaling pathways or populations not captured in our CaMKII guided viral approach, such as non-glutamatergic PBN neurons or thalamic and limbic circuits.

Both CBZ and C194 act predominantly in the periphery at the level of the TG. It is possible that the persistent nature of FRICT-ION causes downstream plasticity that is no longer tightly coupled to peripheral excitability. Earlier timepoints following FRICT-ION may reveal decreased PBN activity in response to systemic analgesics if administered before central plasticity has occurs. Persistent pain also drives alterations in intrinsic and synaptic properties of PBN neurons(60–65), raising the possibility that established central plasticity becomes partially uncoupled from the peripheral mechanisms targeted by C194. Slice electrophysiology will be important for determining whether persistent changes in PBN excitability or synaptic transmission occur after FRICT-ION.

These findings stimulate interest in a broader investigation of the supraspinal circuitry involved in persistent trigeminal pain. The trigeminal system provides parallel projections to the PBN and thalamus, and recent work supports extensive connectivity between the spinal trigeminal nucleus, the PBN, and multiple thalamic subdivisions(66). Future studies should use viral tracing from the TG to define the broader supraspinal network engaged by FRICT-ION and identify neural populations and projection-defined targets recruited during persistent facial pain. Combining tracing, cell-type-specific manipulation, and activity measurements will be particularly important for determining the contributions of thalamic, hypothalamic, and limbic circuits, which may account for behavioral dimensions of facial pain not explained by PBN activity alone.

Together, these findings support the CRMP2-Na_v_1.7 axis as a promising therapeutic target for persistent trigeminal neuropathic pain. At the same time, however, they also highlight the complexity of facial pain. The persistent PBN activity in FRICT-ION mice after C194 treatment despite behavioral improvement suggests that trigeminal pain involves distinct and potentially overlapping adaptations across both peripheral and central circuits. Developing and applying behavioral and circuit level approaches that encompass more aspects of the human pain experience will be essential for determining how peripheral mechanisms, such as the CRMP2-Na_v_1.7 interaction, cooperate with the broader neural adaptations that sustain chronic facial pain.

## Methods

### Animals

#### Sex as a biological variable

All experiments include both male and female animals, and we report similar results for both sexes.

Male and female C57Bl/6J (Jackson labs, #000664) and Na_v_1.7-ΔCRS mice (40) aged 8-20 weeks were used for all experiments. Na_v_1.7-ΔCRS mice have a modified *Scn9a* allele where the 15 amino acid CRMP2 regulatory sequence (CRS) is replaced with a FLAG epitope sequence. Therefore, homozygous Na_v_1.7^ΔCRS/ΔCRS^ mice lack the CRS on both alleles, disrupting regulation of the Na_v_1.7 channel(54). Animals were group housed in a temperature and humidity-controlled room on a 12:12 hour light:dark cycle with *ad libitum* access to food and water.

### Surgery

*FRICT-ION mouse model of trigeminal neuropathic pain.* Surgery was completed as described(38). Anesthesia was induced with 5% isoflurane and maintained at 2.5% isoflurane throughout the procedure. Surgery was performed under a dissecting microscope with the mouse secured in an apparatus equipped with surgical silk restraints that maintained the mouth in an open position. A small incision was made along the right inner buccal mucosa, exposing the maxillary branch of the infraorbital nerve at the foramen rotundum. A 3 mm segment of chromic gut suture (Ethicon #724G, Mettawa, IL, USA) was carefully inserted into the foramen alongside the infraorbital nerve with minimal resistance. The incision was left unsutured, and the mouse was transferred to a heated recovery cage until fully ambulatory. Animals were subsequently monitored daily for body weight, wound healing, and signs of distress or infection. Mice were given 21 days to recover before behavioral testing began. *Viral Delivery and Fiber Optic Cannula Implantation.* Surgery was performed as described(33,51–53,67). Briefly, animals were anesthetized with isoflurane (5% induction, 2.5% maintenance) and secured in a stereotaxic apparatus on a temperature-controlled heating pad. The scalp was shaved and sterilized using 70% ethanol and betadine prior to a midline incision to expose the skull. A craniotomy was performed over the right PBN (A/P −5.15 mm, M/L -1.45 mm, D/V −3.45 mm) and 300 nL of virus carrying a genetically encoded calcium indicator under the calcium calmodulin kinase II promoter (AAV8-CaMKIIα-GCaMP6s-WPRE-SV40, #107790, Addgene, 1.9 x 10^13^ GC/µL, Watertown, MA, USA) was stereotaxically injected into the right PBN using a Nanoject III Auto-Nanoliter Injector (Drummond Scientific, Broomall, PA, USA) at a rate of 1 nL/s, followed by a 5-minute waiting period. Two bone screws were implanted on the left side of midline anterior to bregma and anterior to lambda. A fiber optic cannula (MBF Bioscience LLC, San Diego, CA, USA) was then implanted in the right PBN and secured to the skull with dental cement. Animals recovered from surgery on a heating pad before being returned to their homecage. Mice received a subcutaneous injection of Meloxicam (20 mg/kg, Patterson Veterinary Supply, Alachua, FL, USA) and had access to acetaminophen water (1.1 mg/mL) for 72 hours post surgery(68). Mice were given 21 days to recover before undergoing any behavior experiments. Viral targeting and fiberoptic placement in the PBN were confirmed posthoc via histology and direct visualization of GFP using a Leica Ti2 epifluorescent microscope. No animals were excluded for off target surgeries.

### Drugs

Compound 194 (benzoylated 2-(40piperidinyl)-1,3-benzimidazole analog; 10 mg/kg)(32) and vehicle (10% DMSO, 10% Tween-80, 80% saline) were prepared fresh each day and injected intraperitoneally 1 hour prior to behavioral testing. C194 and vehicle were also administered intranasally (1 mg/mouse, 10 µL/nostril) under light isoflurane anesthesia (∼1.5%) one hour before behavioral testing. Carbamazepine (Sigma-Aldrich, C4024, St. Louis, MO, USA) and vehicle (60 mg/kg in 5% Tween-80, 10% DMSO, 40% PEG-400, 45% saline) were administered via oral gavage 1.5 hours before behavioral testing.

### Behavior

All behavior was performed blinded to pain state (sham v FRICT-ION) and drug injection. All experiments include both male and female mice.

#### Mechanical sensitivity testing

Prior to behavioral testing, mice underwent three days of habituation to acclimate them to the experimental environment, handling, and light restraint. During habituation, mice were individually placed rear-first into loose restraint tubes and placed on a flat surface for 2 hours daily. The tail was secured to the base of the tube with laboratory tape, and a rear stopper was screwed into place to prevent backward movement while allowing the animals’ head and shoulders to remain exposed. Following habituation, mechanical sensitivity was assessed with von Frey filaments applied to the right facial region directly posterior to the whisker pad using the up/down method(69). A positive response was characterized by a rapid facial withdrawal and/or grooming behavior following filament application. A negative response was indicated by the absence of a withdrawal following a 2 second application of the filament bent to a 30° angle.

#### Cold sensitivity testing

The acetone droplet test(70) was applied to the facial region directly posterior to the whisker pad to assess cold hypersensitivity. Directly following the von Frey assay, a 10 µL droplet of acetone was applied to the face, and mice were observed for immediate withdrawal and grooming behavior. The total duration of the response was recorded. Testing was repeated twice per animal, and the average response duration was reported.

#### Open field(70)

Mice were habituated to the room for two days prior to testing. Sixty-decibel white noise was used to mask ambient sounds. Animals were placed in the center of a white Plexiglas open field chamber (45 cm x 45 cm x 40 cm; Stoelting Co, Wood Dale, IL, USA) illuminated by dim light and video recorded for ten minutes. Animals’ movement was analyzed with AnyMaze (Stoelting Co, Wood Dale, IL, USA). The apparatus was cleaned with 70% alcohol after each test between animals.

#### Light-dark preference(71)

Mice were habituated to the room for two days prior to testing. Sixty-decibel white noise was used to mask ambient sounds. The light-dark chamber (Stoelting, San Diego, CA, USA) was placed in the center of the room with a video camera situated directly above to track the animals’ movement. Animals were placed in the light chamber (20 cm x 40 cm) facing away from the entrance to the dark chamber (20 cm x 40 cm) and video recorded for five minutes. AnyMaze was used to analyze animal movement. The apparatus was cleaned with 70% alcohol after each test between animals.

#### Elevated plus maze(72)

Mice were habituated to the room for two days prior to testing. Sixty-decibel white noise was used to mask ambient sounds. The elevated plus maze (Stoelting Co, Wood Dale, IL, USA) was placed in the middle of the room with a video camera situated directly above the apparatus to record the animals’ movement. Animals were placed in the center of the apparatus facing the open arm and video recorded for ten minutes. Animal movement was analyzed with AnyMaze (Stoelting Co, Wood Dale, IL, USA). The apparatus was cleaned with 70% alcohol after each test between animals.

#### Forced swim(73)

Mice were habituated to the room for two days prior to testing. A 2 L beaker filled with room temperature water was placed on a table against a white background. Sixty-decibel white noise was used to mask ambient sounds. A camera supported by a tripod was used to video animal behavior for scoring. The forced swim assay was performed as described(73). Briefly, mice were placed gently in the beaker one at a time and video recorded for six minutes. Following the assay, animals were gently dried off and recovered individually in a cage kept on a heating pad. Immobility time was scored manually using the recorded videos.

*Fiber photometry*. Prior to fiber photometry, mice underwent three days of habituation to acclimate them to the experimental environment, handling, light restraint, and fiber optic patch cord attachment. On the day of testing, animals were habituated in loose restraints with patch cord attached for one hour prior to recording. Calcium transients were continuously recorded (Neurophotometrics FP3002, MBF Bioscience LLC, San Diego, CA, USA) during the stimulation protocol. The right side of the face directly posterior to the whisker pad was stimulated with a 0.07 g von Frey filament, 2.0 g von Frey filament, and 10 µL droplet of acetone and time of application was marked and time-locked in the recording software. Each stimulus was applied three times two minutes apart, and the responses were averaged to represent each animal’s response to that stimuli. GCaMP6s fluorescent signals were analyzed using custom MATLAB scripts. Calcium-dependent signal (470 nm) was normalized to the isobestic control (405 nm) to account for photobleaching and correct for motion artifacts. Changes in fluorescence (ΔF/F) were calculated relative to the average GCaMP6s signal during the 10 seconds directly preceding stimulus application. Area under the curve was calculated for the 5 seconds directly following stimulus application as a measure of stimulus-evoked calcium activity. Fiber photometry was conducted using a within subject design where the same animals were tested before and after FRICT-ION, as well as after drug administration.

*In situ hybridization*. Sham and FRICT-ION animals were deeply anesthetized with 5% isoflurane and transcardially perfused with 20 mL ice cold phosphate buffered saline followed by 20 mL ice cold 10% neutral buffered formalin (NBF). Brains were extracted and post-fixed for 2 hours in 10% NBF at 4°C before being transferred to 30% sucrose at 4°C for 72 hours. Brains were then flash frozen and embedded in optimal cutting temperature gel (OCT, TissueTek) before being sectioned at 20 µm on a cryostat. Three to five representative sections from across the rostro-caudal axis of the PBN were mounted on SuperFrost Plus microscope slides (Fisher Scientific) and allowed to air dry overnight at room temperature. Slides were briefly rinsed in MilliQ water before being immersed in 100% ethanol (2 x 2 minute baths). Tissue was then treated with Protease III for 20 minutes at 40°C in HybEZ oven (ACD Bio). RNAscope was performed as described according to manufacturer’s instructions (ACD Bio)(51,67,74). Briefly, tissue was incubated with the probe for *Fos* (ACD Bio #316921) for two hours at 40°C and then underwent a series of amplification steps (AMP1 30 min, AMP2 30 min, AMP3 15 min) at 40°C. A TSA-based fluorophore was used to develop a fluorescent label before the slides were labeled with DAPI and coverslipped with Vectashield hardset mounting medium (Vector laboratories). Images were acquired on a Nikon Ti2 epifluorescent microscope using a 20x objective. Images were analyzed in QuPath software v0.4.3. The PBN was outlined using the mouse reference atlas (Allen Institute) and DAPI-labeled nuclei surrounded by more than 3 puncta were considered positively labeled *Fos-*expressing cells.

### Statistics

All data was analyzed using GraphPad Prism v11.0.2, using unpaired T-tests, ordinary one-way analysis of variance (ANOVA), repeated measures two-way ANOVAs, and three-way ANOVAs with Tukey’s posttest for multiple comparisons. Statistical significance was determined as P<0.05.

## Study Approval

All procedures were approved by the Institutional Animal Care and Use Committee at the University of Florida, protocol 202400000002.

## Data Availability

All supporting data values are available in the **Supporting data values** document.

## Acknowledgements

We thank Tyler Nelson and Erick Rodriguez for assistance with drug administration and blinding. Schematics were made with BioRender. This work was funded by National Institutes of Health awards F32NS128392 (HNA) and RF1NS131165 (RK) as well as the Facial Pain Research Foundation (RK). Generation of the Na_V_1.7-ΔCRS mouse line was made possible by a pilot grant from the Genetically Engineered Mouse Models (GEMM) Core at the University of Arizona (RK). We thank Dr. Teodora G. Georgieva for generating the mouse line.

## Author contributions

HNA and RK developed the concept. HNA and NKG designed and conducted the experiments, completed the data analysis, and wrote the manuscript. All authors had the opportunity to discuss the results and comment on the paper.

